# High-resolution cross-species transcriptomic atlas of dorsal root ganglia reveals species-specific programs for sensory function

**DOI:** 10.1101/2022.06.21.497049

**Authors:** Min Jung, Michelle Dourado, James Maksymetz, Amanda Jacobson, Miriam Baca, Oded Foreman, David H. Hackos, Lorena Riol-Blanco, Joshua S. Kaminker

**Author notes:** indicates equal contribution (Min Jung, Michelle Dourado).

## Abstract

Sensory neurons of the dorsal root ganglion (DRG) play a crucial role in maintaining tissue homeostasis by sensing and initiating responses to stimuli. While most preclinical studies of DRGs are conducted in rodents, much less is known about the mechanisms of sensory perception in primates. We generated a transcriptome atlas of mouse, guinea pig, cynomolgus monkey, and human DRGs using a framework that implements a common laboratory workflow and multiple data-integration approaches to generate high-resolution cross-species mappings of sensory neuron subtypes. Using our atlas, we identified conserved core modules highlighting subtype-specific biological processes related to inflammatory response. We also identified divergent expression of key genes involved in DRG function, suggesting species-specific adaptations. Among these, we validated that Tafa4, a member of the druggable genome, was expressed in distinct populations of DRG neurons across species, highlighting species-specific programs that are critical for therapeutic development.

## INTRODUCTION

The dorsal root ganglion (DRG) plays a key role in the perception of sensory information. A primary function of sensory neurons within the DRG is the detection of stimuli such as touch, noxious stimuli, temperature, itch or proprioception, and to transmit this information to the central nervous system ^1–3^. This ability to precisely differentiate and process distinct sensory information is in part enabled by the extensive heterogeneity of sensory neuron subtypes which can differ in cell body size, degree of myelination, conduction velocities, and innervation tissue4,5.

Rodents have been the primary model system used to study somatosensory function. Unbiased descriptions of the rodent sensory neurons using single-cell RNA sequencing have provided opportunities to map transcriptional traits to anatomical and functional properties ^6–8^. From these studies, rodent DRG neurons have been moleculary classified into different groups: heavily myelinated limb proprioceptors and A-fiber low-threshold mechanoreceptors (LTMRs) that express neurotrophin receptor tyrosine kinases (Ntrk2 and Ntrk3); C-fiber LTMRs that express tyrosine hydroxylase (Th) and Vglut3 (Slc17a8); C-fiber non-peptidergic nociceptors marked by the expression of Mrgprd, Mrgpra3, and Sst; non-myelinated C-fiber and lightly myelinated A*δ*- fiber peptidergic nociceptors which express a variety of neuropeptides including substance P (Tac1), calcitonin related peptide (Calca) and pituitary adenylate-cyclase-activating neuropeptide (Adcyap1) along with Ntrk1; A*δ*-fiber and C-fiber nociceptors that express the cooling and menthol sensing receptor, TrpM8. In rodent models, these markers correlate well with previous electrophysiological and neurochemical characterization, and have enabled us to broaden our basic understanding of somatosensation.

However, even with our extensive understanding of rodent sensory neurons, the translation of somatosensory mechanisms from preclinical models to the clinic remains challenging due to molecular differences between sensory neurons present in rodents and humans. For example, a considerable subpopulation of neurons that co-express NTRK1 and RET were observed in human DRGs, but were absent from mouse DRGs ^9^. Additionally, numerous studies have highlighted that Nav1.8, Nav1.9, P2X3 receptor and TRPV1 are essentially present in all human DRG neurons ^10–12^, whereas in mice these genes are expressed in specific sub-populations of DRG neurons ^8, 13^. These differences between species accentuate the need for comprehensive, cross-species molecular studies of sensory neurons within the DRG.

Recent transcriptome studies have enabled deeper understanding of the molecular landscape of primate DRGs^14–16^. However, some of these studies suffer from hard-to-interpret sensory neuron subtype mapping due to differences in sequencing technologies, laboratory protocols, or sample archival methods. These differences introduce technical artifacts that make it challenging to precisely integrate such transcriptome data from different species, which is crucial for comparison of transcriptional programs across pre-clinical and clinical specimens. Various computational methods have been applied to align homologous cell types between species while attempting to address technical artifacts. These include Seurat for aggregating scRNA-seq datasets using anchors identified from canonical correlation analysis ^17^ and MetaNeighbor for quantifying cell-type replicability across data sets ^18^. While studies have demonstrated the utility of these methods in cross-species cell-type homology mapping ^19–21^, interpretation and evaluation of their performance requires significant computational and biological expertise.

Ultimately, appropriately pairing laboratory protocols and technologies with the most relevant computational methods is of utmost importance for the creation of meaningful, high-resolution cross-species mappings at a single-cell resolution. Mappings between preclinical models and humans are fundamental for characterizing gene expression similarities and differences between species to inform relevant therapeutic hypotheses.

Numerous studies have highlighted the therapeutic potential for targeting the neuro-immune axis ^22–30^. For example, Hoeffel et. al. has shown that in mice TAFA4, a chemokine-like protein which is secreted by sensory neurons, shifts dermal macrophages toward a phenotype that promotes tissue healing. And, *Kambrun et al* and *Yoo et al.* showed that in mice C-LTMRs secrete TAFA4, which promotes microglial process retraction and results in decreased production of microglial mediators known to alleviate mechanical pain hypersensitivity. As Tafa4 is being evaluated for use in a clinical setting^28^, it is crucial to determine which neuronal populations in humans express TAFA4 and how these map to their preclinical counterparts – such data would inform therapeutic development related to this signaling molecule. In this paper, we (1) describe a protocol for efficient isolation of DRG nuclei from multiple species, (2) provide the first high-resolution, comprehensive, detailed single-nucleus transcriptome atlas of DRG from pre-clinical to human samples, and (3) characterize the transcriptional convergence and divergence of sensory neuron subtypes from rodents to humans. Our results reveal that DRG sensory neuron subtypes are in general well-conserved across species. However, we identified key differences in gene products involved in pathophysiological processes which point to the potential for species-specific sensory neuron functions. Understanding the molecular and functional similarities and differences between somatosensory neurons in rodents and primates will enable a better understanding of the role of these neurons in sensory perception and tissue homeostasis, facilitating therapeutic efforts targeting sensory neurons.

## RESULTS

### Fresh tissue and density-gradient centrifugation enable DRG neuron enrichment in single-nucleus RNA-seq

To characterize the transcriptome of preclinical and clinical DRG samples, we developed a protocol to isolate nuclei from fresh or frozen samples, and generated transcriptome data from DRGs from multiple species. We harvested DRGs from 5 mice, 2 guinea pigs, 3 cynomolgus monkeys and 7 human donors (Fig. 1a; Supplementary Table 1). Distribution of sex and vertebral level varied across samples and species (Supplementary Table 1). For all preclinical species, both fresh and frozen tissues were collected, whereas only frozen samples were obtained for human specimens (Fig. 1a; Supplementary Table 1). From these DRGs we isolated nuclei using two different protocols: a FACS-based method, and a density-gradient (DG) centrifugation method (Fig. 1a). Analysis of nuclei isolated from the FACS protocol revealed that these samples contained fewer larger nuclei, which could suggest a loss of neuronal nuclei during the sorting process (Fig. 1b). The large size of the DRG sensory neuron nuclei could make them more susceptible to the shear forces within the sorting nozzle, compromising their nuclear membranes. Single nuclei cDNA libraries were prepared using a droplet-based system from the 10X Genomics platform (see “Methods”). Computational analysis of the single-nuclei RNAseq data was performed using a standard analysis pipeline (Seurat V3; see “Methods”). We removed low-quality nuclei based on quality metrics including number of transcripts, number of genes, and number of mitochondrial reads. We then merged data generated from different isolation methods and tissue types (i.e., fresh and frozen) using the anchoring-based integration approach, and clustered the cells using the graph-based clustering approach implemented in Seurat (see “Methods”; Fig. 1a).

**Fig. 1.**
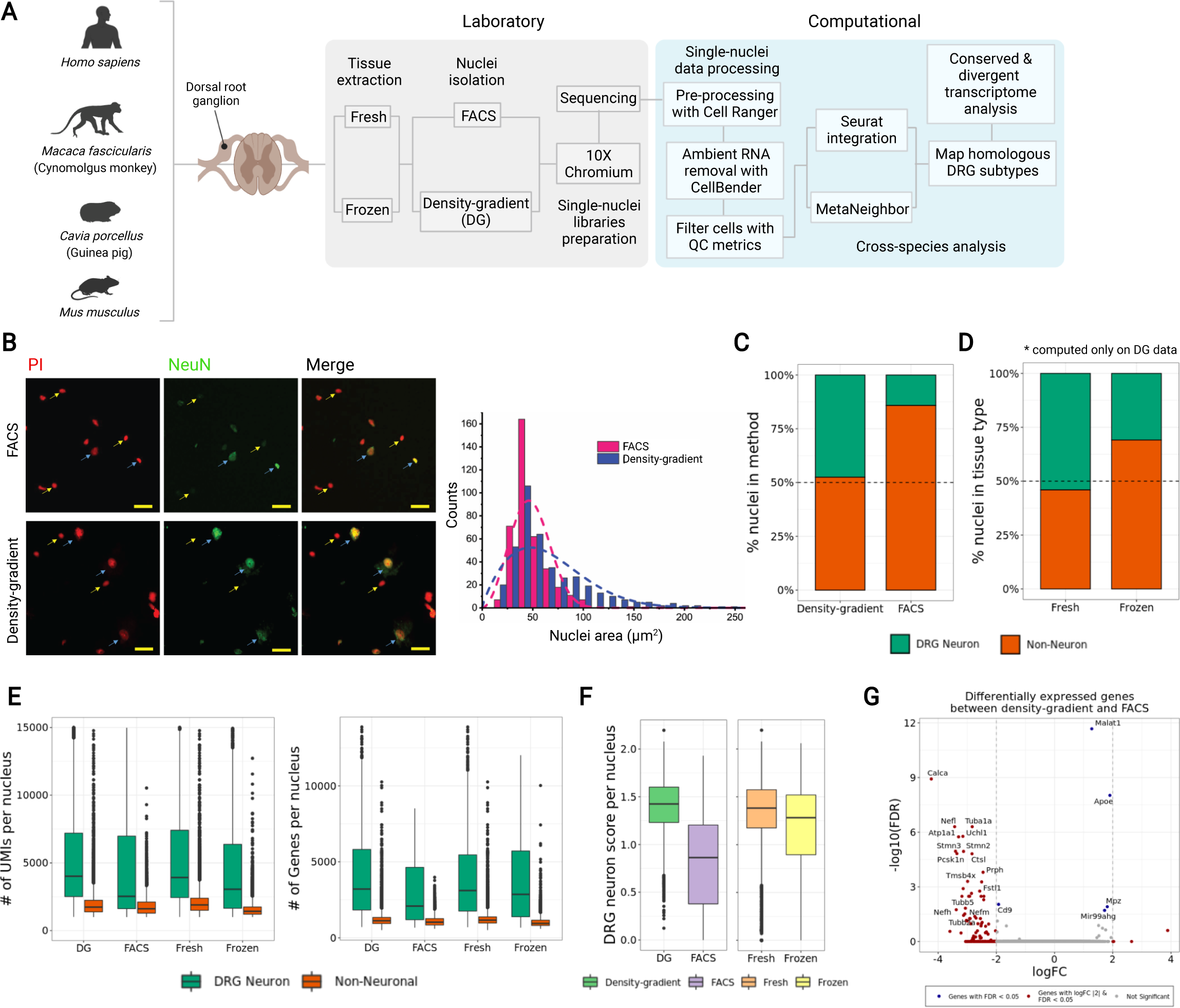
Comparison of nuclei isolation methods and tissue types in dorsal root ganglia single-nucleus RNA-seq. **a** Overview of laboratory and computational workflow. **b** Micrographs of nuclei from FACS and density-gradient protocols. Blue arrows indicate neuronal nuclei (PI+ and NeuN+) and yellow arrows indicate non-neuronal nuclei (PI+ and NeuN-). Distribution of nuclei size by capture method. Dotted lines represent smoothed distribution of binned data. c-g Summary metrics computed for different nuclei isolation methods and tissue types in mouse data. **c-d** Relative cell-type composition for the nuclei isolation methods or tissue types. **e** Number of transcripts and genes detected per nucleus for different nuclei isolation methods and tissue types. **f** DRG neuron signature score for different nuclei isolation methods and tissue types. **g** Volcano plot showing results from differential expression analysis of transcripts from FACS versus DG nuclei. Statistically significant genes are indicated by red or blue colors.

Comparison of relative cell-type composition between the two isolation methods from mouse DRG samples revealed that the DG protocol captured more DRG neurons than the FACS method (Fig. 1c and Supplementary Fig. 1a). Similar differences were seen in data generated from other species (Supplementary Fig. 1a). We also observed a similar enrichment of DRG neurons in data from both fresh and frozen tissue samples (Fig. 1d and Supplementary Fig. 1b). The nuclei captured from mouse DRG samples contained 2,062 genes and 6,489 unique transcripts per nucleus, on average. Regardless of the nuclei isolation method or the type of tissue sample (fresh or frozen), we detected more transcripts and genes in neurons than non-neuronal nuclei (Fig. 1e). To further characterize the neurons from our single nuclei data sets, we computed a DRG neuron signature score, reflecting the mean expression levels of DRG-specific marker genes curated from previous reports ^7, 14, 16^. The median DRG neuron signature score per nucleus was higher in DG than FACS data, whereas the median DRG signature scores between fresh and frozen tissues were comparable (Fig. 1f). Finally, to determine if there were protocol-specific transcriptional differences, we performed differential expression analysis between DG and FACS data. Only 0.2% (41 out of 18545) of expressed genes were statistically differentially expressed between two methods (Fig. 1g). Within the small number of genes that were differentially expressed, we noticed a slight enrichment of cell type-specific genes that could reflect cell compositional differences resulting from the FACS and DG methods (Fig. 1b, c). In general, the FACS and DG protocols produced broadly similar transcription profiles.

Together, these data suggest that the DG nuclei isolation protocol improved our ability to capture and generate the transcriptome data of sensory neurons from either fresh or frozen tissue samples, and provide a framework for isolation and characterization of DRG nuclei from other species.

### snRNA-seq identifies 17 distinct sensor neuron subtypes in the mouse DRG, including characterization of a unique subpopulation of C-LTMRs

We integrated expression data from 37,384 nuclei from 5 mouse DRGs (Supplementary Table 1). After clustering, we annotated each cluster with canonical markers (Fig. 2a; Supplementary Table 1). Sensory neurons were identified by their expression of Snap25, Rbfox3, Scn10a, Nefh and Calca; clusters expressing Apoe and Fabp7 were identified as satellite glia (SG); myelinating Schwann cell (mSC) clusters were identified based on their expression of Mpz, Mbp, and Plp1; macrophage (MP) clusters were identified by their expression of Lyz2 and Csf1r; Pecam1 and Vwf were used to identify clusters of endothelial cells (EC); and Dcn, Pdgfra and Fn1 were used to identify fibroblasts (FB) clusters.

**Fig. 2.**
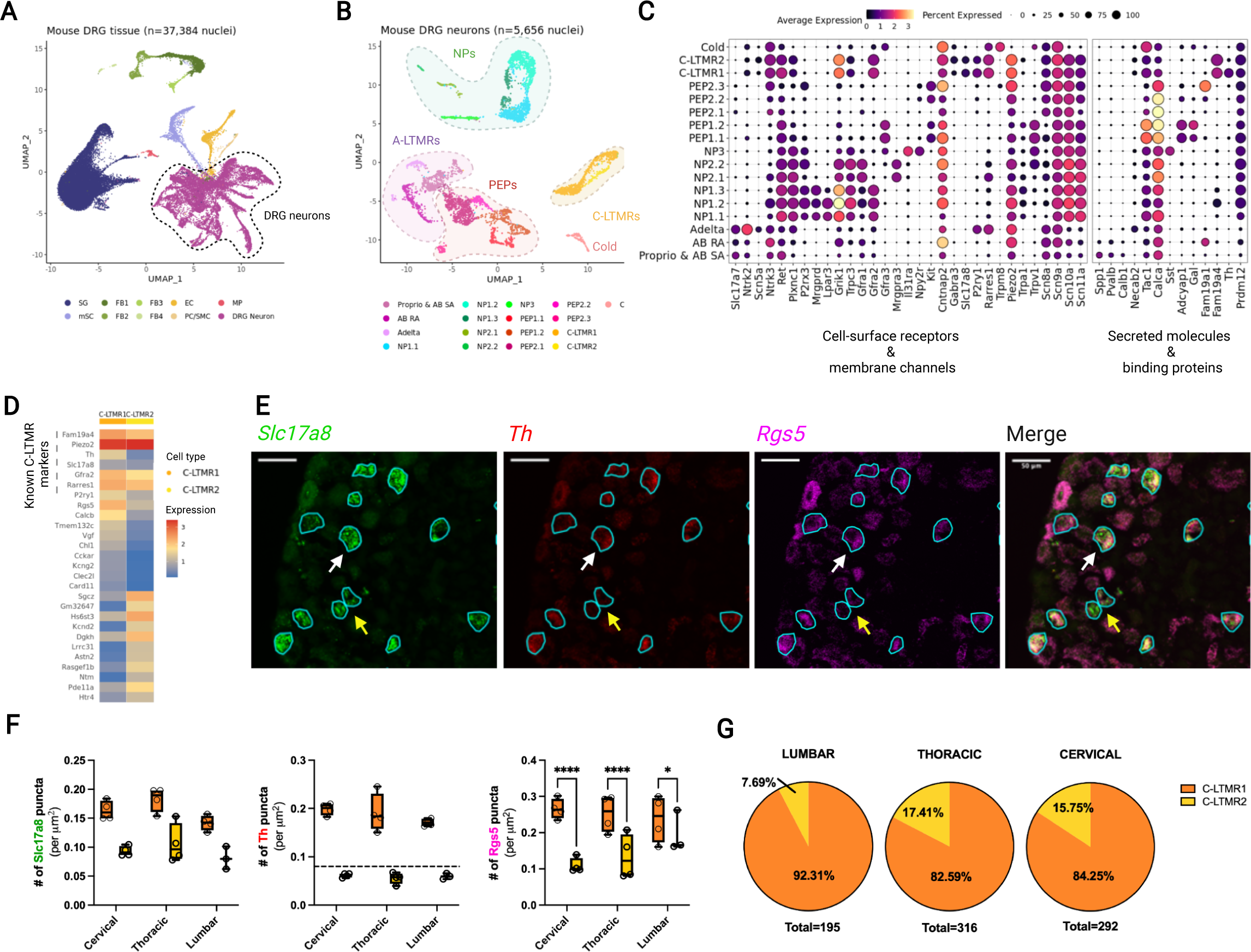
Single-nuclei transcriptome atlas of mouse dorsal root ganglia (DRG) sensory neurons. **a** UMAP of 37,384 mouse DRG nuclei colored by cell types and annotated by marker genes as indicated in the main text. **b** UMAP of 5,656 mouse DRG neurons colored by subtypes and annotated by marker genes as indicated in the main text. **c** Fraction of nuclei (dot size) in each subset expressing canonical marker genes (columns) and their scaled average expression level in expressing cells (dot color). **d** Heatmap comparing mean expression levels (color bars) of top differentially expressed genes (rows) between C-LTMR1 and C-LTMR2 subtypes. **e** RNAScope validation of Rgs5, Th and Slc17a8 expression in mouse DRGs. White arrows indicate Slc17a8-positive cells expressing both high levels of Th and Rgs5 and yellow arrows indicate cells that express Slc17a8-positive cells but low amounts of Th and Rgs5 transcript. **f** Quantification of RNA transcript punctate dots (representing expression level) normalized by slide area for each DRG level and C-LTMR subtype **g** Distribution of the two C-LTMRs subpopulations by DRG level.

To provide better resolution on the sensory neuron population, we removed clusters containing non-neuronal nuclei, and performed iterative clustering on the remaining presumptive neuronal nuclei. Additionally, we filtered any nuclei expressing satellite glia-specific transcripts (Plp1, Sparc, Mpz). From this filtering, we retained 5,656 mouse DRG nuclei and annotated each cluster using known mouse DRG sensory neuron subtype-specific markers (Fig. 2b; see also ^7, 8, 14–16^. We identified 17 transcriptionally distinct DRG sensory neuron subtypes that differ in expression of both cell-surface and secreted molecules (Fig. 2b,c). Large diameter, fast conducting myelinated A-fibers that express Nefh include: proprioceptors which express Pvalb and Spp1; A*β* SA-LTMRs which express Calb1; A*β* RA-LTMRs which highly express Ntrk3; and A*δ* LTMRs which express Necab2 and Scn5a. Non-peptidergic C-fiber nociceptive neurons which express Trpc3 include: non-peptidergic nociceptor type1 (NP1.1, 1.2 and 1.3) which are marked by expression of Mrgprd, Gfra1, and Gfra2; non-peptidergic nociceptor type2 (NP2.1 and 2.2) which express Mrgpra3 and Gfra1 but not Gfra2; and non-peptidergic nociceptor type3 (NP3) which express pruritogen related genes like Sst, Il31ra, and Nppb. Peptidergic C-fiber nociceptors which express Calca include: peptidergic nociceptor type 1 (PEP1.1 and 1.2) which express a variety of neuropeptides including Tac1, Adcyap1, Gal, and Trpa1; and peptidergic nociceptor type 2 (PEP2.1, 2.2, and 2.3) which express Kit, Fam19a1 and Cntnap2. Cold thermoreceptor neurons distinctly express Trpm8 but not Piezo2. Finally, unmyelinated C-fiber LTMRs express Th, Slc17a8, Fam19a4, Gfra2 and Piezo2.

Within our mouse data, we identified an additional cluster of C-LTMR neurons (Fig. 2b,c). The two C-LTMR clusters (C-LTMR1 and C-LTMR2) express Slc17a8 (Vglut3), but differ in expression intensity of other known C-LTMR specific markers including Th, Fam19a4, Rarres1, P2ry1, and Gfra2 (Fig. 2d). *Renthal et al* also noted two C-LTMR subtypes that differ in Th expression ^7^. We performed differential expression analysis of the two C-LTMR populations (Fig. 2d), and identified that genes more highly expressed in C-LTMR2 are associated with signaling pathways and axon guidance gene ontology terms (Supplementary Fig. 2a).

Additionally, we performed *in situ* hybridization (ISH) validation on C-LTMR markers predicted by differential expression analysis to distinguish C-LTMR1 from C-LTMR2 neurons (Fig. 2e). Consistent with our transcriptome data, our *in situ* data reveal that C-LTMRs could be divided into two groups with differing Th expression, regardless of vertebral level, highlighting the utility of this marker to subset these two subtypes (Fig. 2e,f). Additionally, expression of Rgs5, a regulator of the G-protein signaling family, is correlated with high expression of Th in C-LTMR1 neurons, regardless of the level of the DRG. While we did not observe any differences in the cell diameter size of these two populations of C-LTMRs (Supplementary Fig. 2b), we did find that the proportion of C-LTMR subtypes differed by DRG levels, with lumbar having the fewest C-LTMR2 (7.69% in lumbar versus 17.41% in thoracic and 15.75% in cervical; 4 biological replicates per level; Fig. 2g).

Overall, our single-nuclei expression data identified the major known sensory neuron populations in the mouse DRG. Additionally, a second subpopulation of C-LTMR sensory neurons were identified that differ both in their transcriptional profiles and also their proportion across DRG levels, which could reflect that these neurons transduce distinct sensory information.

### Cross-species mapping of DRG sensory neuron subtypes across mouse, guinea pig, cynomolgus monkey, and human

To further expand our understanding of somatosensory function in higher species, we performed single nucleus RNAseq analysis of DRGs from guinea pig, cynomolgus monkey, and human samples, from 2 donors, 3 donors, and 7 donors, respectively (Fig. 3a; Supplementary Table 1). We profiled 31,776 nuclei from guinea pig DRGs, 103,383 nuclei from cynomolgus monkey DRGs, and 111,925 nuclei from human DRGs (Supplementary Fig. 3a). H&E imaging of the tissues suggests that while the neuron size increases from mouse to cynomolgus monkey to human, we did not notice an appreciable difference in the size of satellite glia (Supplementary Fig. 3b). The same analysis pipeline used for analysis of the mouse transcriptome data was used for these species-specific datasets (see “Methods”).

**Fig. 3.**
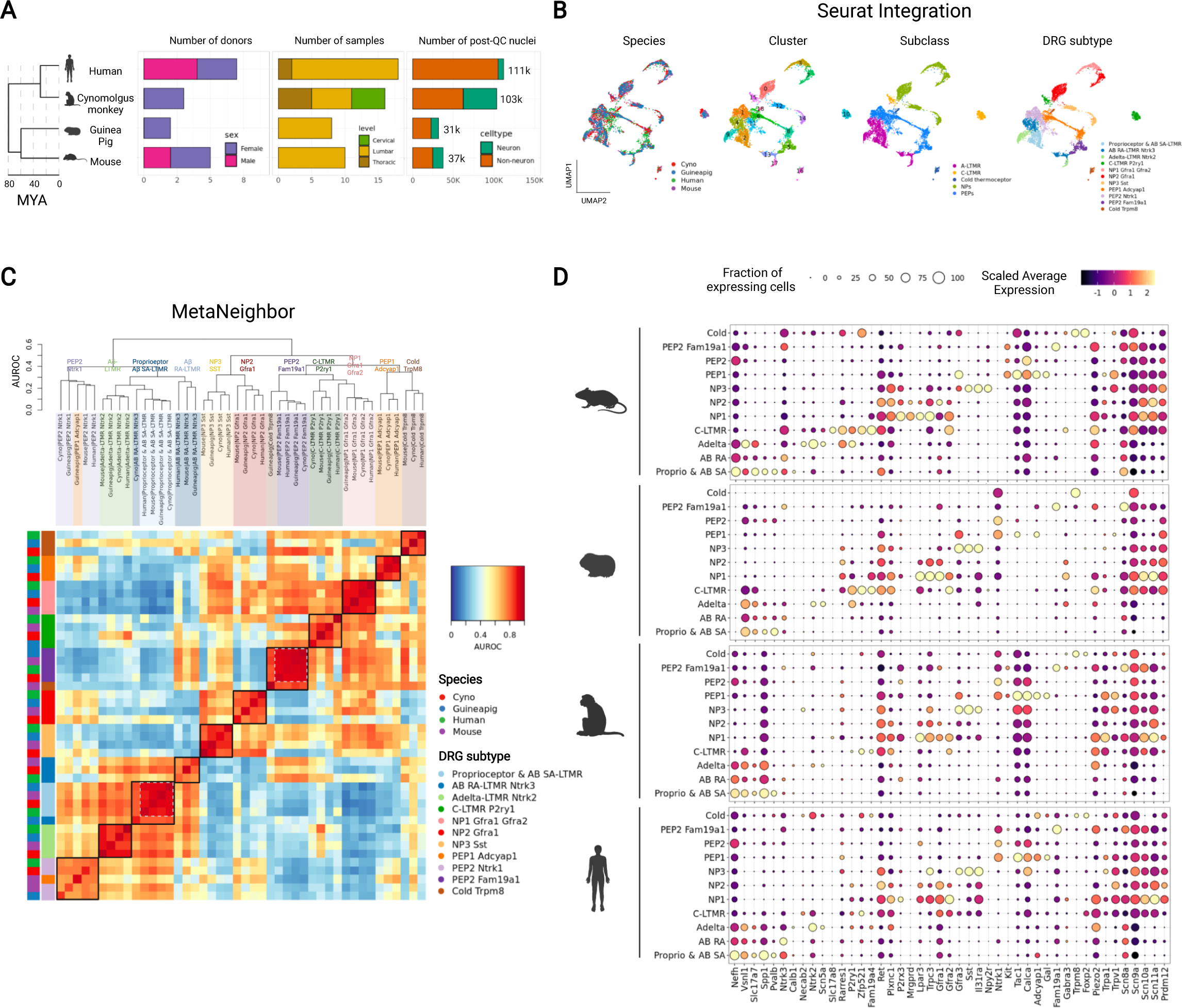
DRG sensory neuron homology consensus across mouse, guinea pig, cynomolgus monkey and human. **a** Summary metrics for mouse, guinea pig, cynomolgus monkey, and human DRG single-nuclei RNA-seq data. Evolutionary distances (Million Years Ago; MYA) are shown in dendrogram to the left. **b** UMAP of Seurat-integrated DRG neurons across mouse, guinea pig, cynomolgus monkey, and human, faceted and colored by species, Seurat clusters, DRG subclasses, and subtypes. **c** Cross-species dendrogram and heatmap of MetaNeighbor AUROC scores (y-axis in plot) colored by DRG sensory neuron subclasses. **d** Fraction of nuclei (dot size) in each neuron subtype expressing selected canonical markers (columns) and their scaled average expression level in expressing nuclei (dot color) in mouse, guinea pig, cynomolgus monkey and human data.

To map DRG sensory neuron populations across species, we performed comparative analysis using two different methods, Seurat and MetaNeighbor (see “Methods”). We used Seurat for aggregating our cross-species datasets using anchors identified from canonical correlation analysis ^17^ and MetaNeighbor for quantifying cell-type replicability in cross-species datasets ^18^. Previous cross-species studies ^19–21^ demonstrated that both Seurat and MetaNeighbor provide effective computational methods to map cell types across independent datasets to reveal cell-type relationships among species. Using Seurat, we integrated DRG sensory neuron subsets across species (Fig. 3b). Sensory neuron nuclei were well-integrated between different species and clustered according to their subtype identities, rather than by species. (Fig. 3b). In parallel to our approach using Seurat, we also performed MetaNeighbor analysis on the DRG neuron subsets. The dendrogram of the mean area under the receiver operator characteristic (AUROC) curve scores was organized by the major DRG subtype category rather than by species (Fig. 3c). Together, the results of these analyses, derived from two distinct computational approaches, are well-aligned (Fig. 3d), highlighting that sensory neuron subtypes are generally well-conserved across species. Our computational strategy enabled cross-species, high-resolution DRG cell type annotation, providing interpretable comparison of gene expression profiles from preclinical models to humans.

Leveraging our well-aligned cross-species data, we generated a dot plot to compare the expression patterns across species of genes known for their roles in sensory function (Fig. 3d). These data highlight known gene expression profiles, such as the ubiquitous expression of Scn9a (Nav1.7) across all sensory neuron subtypes. Our data also show that Scn8a (Nav1.8) is primarily expressed in NPs, PEPs and C-LTMRs across all species. Additionally, our data highlight a number of other cell-type specific expression patterns. For example, proprioceptors and A*β* SA-LTMRs express known markers including Pvalb, Ntrk2, Ntrk3 and Slc17a7; A*β* RA-LTMRs express high levels of Ntrk3, Slc17a7, Vsnl1; and A*δ*-LTMRs distinctively express Ntrk2 and Scn5A; C-LTMRs express Gfra2, Piezo2 and P2ry1 along with the newly identified C-LTMR-specific marker^16^, Zpf521/ZNF521; NP1 express Gfra1, Gfra2, Trpc3, and Plxnc1; NP2 express Gfra1, Trpc3 and Plxnc1, but are negative for Gfra2; and NP3 express Sst and Il31ra; PEP1 express Gal, Adcyap1, and Trpa1; and PEP2 Ntrk1 express Ntrk1, Tac1, Calca, and Nefh; PEP2 Fam19a1 express Fam19a1, Piezo2, and Kit; finally, cold thermoreceptors express Trpm8, Foxp2 but do not express Piezo2.

We also compared our data to other recently generated primate expression data ^14–16^ and found that, in general, markers identified within these other analyses map to similarly described DRG subtypes within our data (Supplementary Fig. 4a). For example, markers used in Kupari et al to classify DRG subtypes from rhesus macaque, were expressed in similarly identified populations of our cynomolgus monkey data except for A-LTMRs where our data captured different subpopulations of the A-LTMRs (i.e Proprioceptor, A*β* SA-LTMRs, A*β* RA-LTMRs, and A*δ*- LTMRs). When we compared our human data with recent single-nuclei RNA-seq data from Nguyen et al. and spatial transcriptome data from Tavares-Ferreira et al., we observed differences in subtype classification across these data sets that could be a result of cluster resolution driven by either technological or analytical methods. While these studies provide additional data for understanding the transcriptional landscape of human DRGs, the inconsistent DRG subtype nomenclature from mouse to human makes it challenging to compare and interpret gene expression profiles from these datasets to those generated from preclinical models. The data on their own cannot be used off-the-shelf, but as we have shown, require careful integrative analysis that takes into account technical or biological differences between datasets. Our approach produced a well-integrated, cross-species atlas with harmonized DRG subtype classification that enables comparative analysis of sensory neurons.

### Conserved and species-specific transcriptional programs in DRG sensory neurons

Leveraging the integrated single nuclei data, we performed analyses to assess the similarities and differences of sensory neurons across species. First, we compared the relative proportions of sensory neurons across species by calculating the frequency of each subtype within individual species (analysis restricted to lumbar level DRGs; Fig. 4a). We observed a smaller proportion of C-LTMRs in cynomolgus monkey and human (∼1% and ∼0.8% respectively), as compared to guinea pig and mouse (8% and 14%, respectively). We also noted a larger proportion of NP1 and NP2 subtypes in cynomolgus monkey as compared to other species. Additionally, the proportion of Trpm8 neurons in guinea pig and cynomolgus monkey was far lower than in other species, and the proportion of PEP2 Fam19a1 sensory neurons was lower in cynomolgus monkey as compared to other species. Notably, we observed that peptidergic neurons (PEP1, PEP2 Ntrk1 and PEP2 Fam19a1) comprise a majority of the sensory neuron populations from human DRGs within our analysis.

**Fig. 4.**
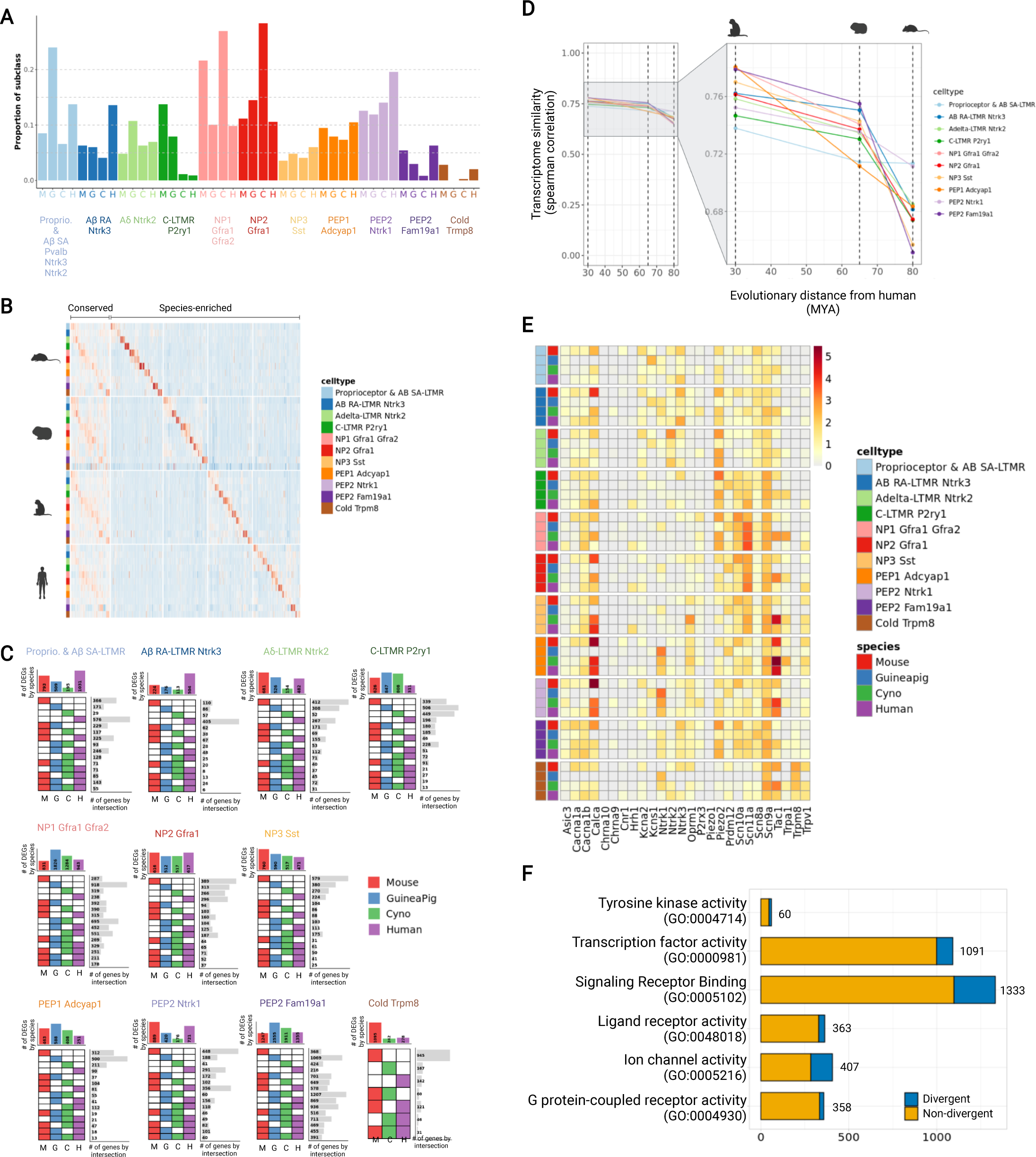
Conserved and species-specific transcriptional programs in DRG sensory neurons. **a** Relative DRG subtype proportions across all species. **b** Customized UpSet plot showing number of conserved and species-specific genes across datasets for each of the major sensory neuron subtypes, as indicated above each plot. Total number of genes that are unique to one species (one colored box per row), or shared across multiple species (multiple colored boxes per row), are indicated to the right of each row. **c** Heatmap of conserved genes and species-specific genes by DRG subtypes from panel (b). **d** Scatter plots of transcriptome correlation (Spearman) of DRG subtypes between human and other species. **e** Heatmap indicating normalized expression levels of genes associated with pain perception in each neuronal subtype. f Number of genes with divergent or non-divergent subtype specific expression between mouse and human DRGs indicated by gene ontology(GO) annotation category. Numeric numbers on each bar represent the total number of genes in a given GO category.

Next, to characterize core programs of sensory function, we performed differential expression analysis to identify subtype specific gene expression programs within each species. Then we took the intersection of these gene lists to collect those subtype specific genes that were common across multiple species (Fig. 4b,c; Supplementary Table 2 for complete list of genes; see “Methods”). For subtype-specific genes conserved across all four species, we performed gene ontology analysis to assess conserved biological programs (Supplementary Fig. 5a). Notably, this analysis revealed enrichment of distinct biological pathways across sensory neuron subtypes which reflect that a variety of cellular pathways are utilized differentially by distinct subtypes to transduce sensory information. For example, A*β* RA-LTMR subtype, which is identified by expression of Ntrk3, Scn1a, and Atp2b2, is enriched for genes associated with *action potential conduction and rapid response to mechanical stimuli*, and could point to biological programs most relevant for sensing differences in touch sensation ^31^. Additionally, the PEP1 subtype, which is identified by expression of Adcyap1, Trpv1, Tac1 and Oprm1, is enriched for genes associated with *acute inflammatory response* and the perception and modulation of *pain signaling.* The PEP1 subtype could be useful in identification of conserved neuro-immune signaling pathways, which might also be relevant for therapeutic development.

We also examined correlations between the transcriptomes of species pairs to gain further insight into the evolutionary divergence of sensory neuron subtypes. Here, for each species we determined the average expression of individual genes by subtype. Then, between each pair of species and for each subtype, we calculated the correlation of average gene expression across all genes. To further assess the relationships of preclinical models to humans, we plotted the transcriptome correlation by subtype for each species in relation to human (transcriptome correlation metrics between each species and human are indicated by the dots in Fig. 4d). This highlighted that cynomolgus monkey and human share the highest correlation between transcriptomes across all subtypes. Our analysis further revealed that transcriptomes of Proprioceptor & A*β* SA-LTMRs are the most correlated between human and mouse (R = 0.71), while the transcriptomes of the NP1 and PEP2 Fam19a1 subtypes (R=0.65 and 0.65, respectively) are the least correlated between mouse and human (Supplementary Fig. 5b). In general, the transcriptome of each sensory neuron subtype becomes less correlated as the evolutionary distance from human species increases (Fig. 4d). Finally, we examined the expression of a set of genes including ion channels, ligand-gated channels, G-protein coupled receptors (GPCRs), and neuropeptides that have been well-studied for a role well-studied for a role in sensory transduction and could provide therapeutic targets (Fig. 4e, Supplementary Fig. 6-7). From this analysis, we observed divergent expression patterns of these sensory molecular determinants across species. For example, analysis of genes involved in pain perception (Fig. 4e) highlighted that Trpv1 was expressed in more broader sets of subtypes including C-LTMRs, all NPs, all PEPs and cold thermoreceptors in human DRGs, while Trpv1 expression was restricted to subsets of NPs, PEPs, and cold thermoreceptors in mouse, guinea pig and cynomolgus monkey DRGs. Additionally, Scn8a was expressed in essentially all DRG subtypes in primates whereas Scn8a was selectively expressed in LTMRs and PEPs in mouse.

Leveraging these data from our harmonized cross-species atlas enabled detailed interrogation of individual genes and gene sets from preclinical models to human, and pointed to divergent and conserved biological processes. Further, these data can help inform therapeutic programs targeting sensory neuron-mediated pathophysiological processes such as pain perception, and can also help inform our understanding of any species-specific adaptations that have evolved.

### TAFA4 displays divergent expression pattern from rodents to human

We were interested in performing a broad survey of the expression of genes associated with the druggable genome. For this we collected the list of genes annotated with the ontology terms ion channels, G protein-coupled receptor, tyrosine kinase, ligands, signaling-pathway related molecules. We assessed the expression of these genes across preclinical and human samples and identified those genes that displayed divergent expression patterns between mouse and humans (Fig. 4f). Interestingly, within the “Signaling Receptor Binding” category we observed that 18% (235) of the 1,333 genes were expressed in distinct subtypes of DRG neurons between mouse and human.

One gene of interest from the “Signaling Receptor Binding” category is TAFA4 (Fam19a4) which is expressed by sensory neurons and has been shown to play a role in maintenance of tissue homeostasis by modulating the function of Il10+ dermal macrophages and microglia ^27, 28, 30^. In our mouse data, TAFA4 is predominantly expressed by C-LTMRs and is also expressed by some NPs. Similarly in guinea pig and cynomolgus monkey, TAFA4 is expressed in both C-LTMRs and NPs and (Fig. 5a). Notably, the expression of TAFA4 in our human data is distinct from the preclinical models as TAFA4 is most strongly expressed in A*δ*-LTMRs that co-express NTRK2 and SCN5A, and is also expressed in cold sensing neurons and C-LTMRs. To validate these findings in tissue samples, we performed *in situ* hybridization (ISH) to assess the expression of TAFA4 transcripts in NTRK2+ neurons. Consistent with our single nuclei RNAseq data, FISH images revealed that ∼98% of TAFA4+ neurons are NTRK2+ in human DRGs whereas in DRGs from other species, we observed very few cells with expression of both markers (∼9% in mouse, ∼1.3% in GP and ∼9% in Cyno.) (Fig. 5b,d). A*δ* fibers are characterized as having a larger cell body diameter than C fibers. Therefore, to determine if the expression of NTRK2 and TAFA4 was observed more frequently in larger diameter neurons in human samples as would be predicted from our data, we assessed the average cell body area of neurons across all species that express TAFA4 and NTRK2. Consistent with our single nuclei RNAseq and *in situ* data we observed that the cell area of TAFA4+ neurons in mouse, guinea pig and cynomolgus monkey predominantly fell into small-to-medium range whereas in human samples, this cell area distribution shifts towards larger neurons (Fig. 5c). Together these data indicate significant differences in the cell-type expression of TAFA4 across species, which will have implications for therapeutic strategies targeting TAFA4.

**Fig. 5.**
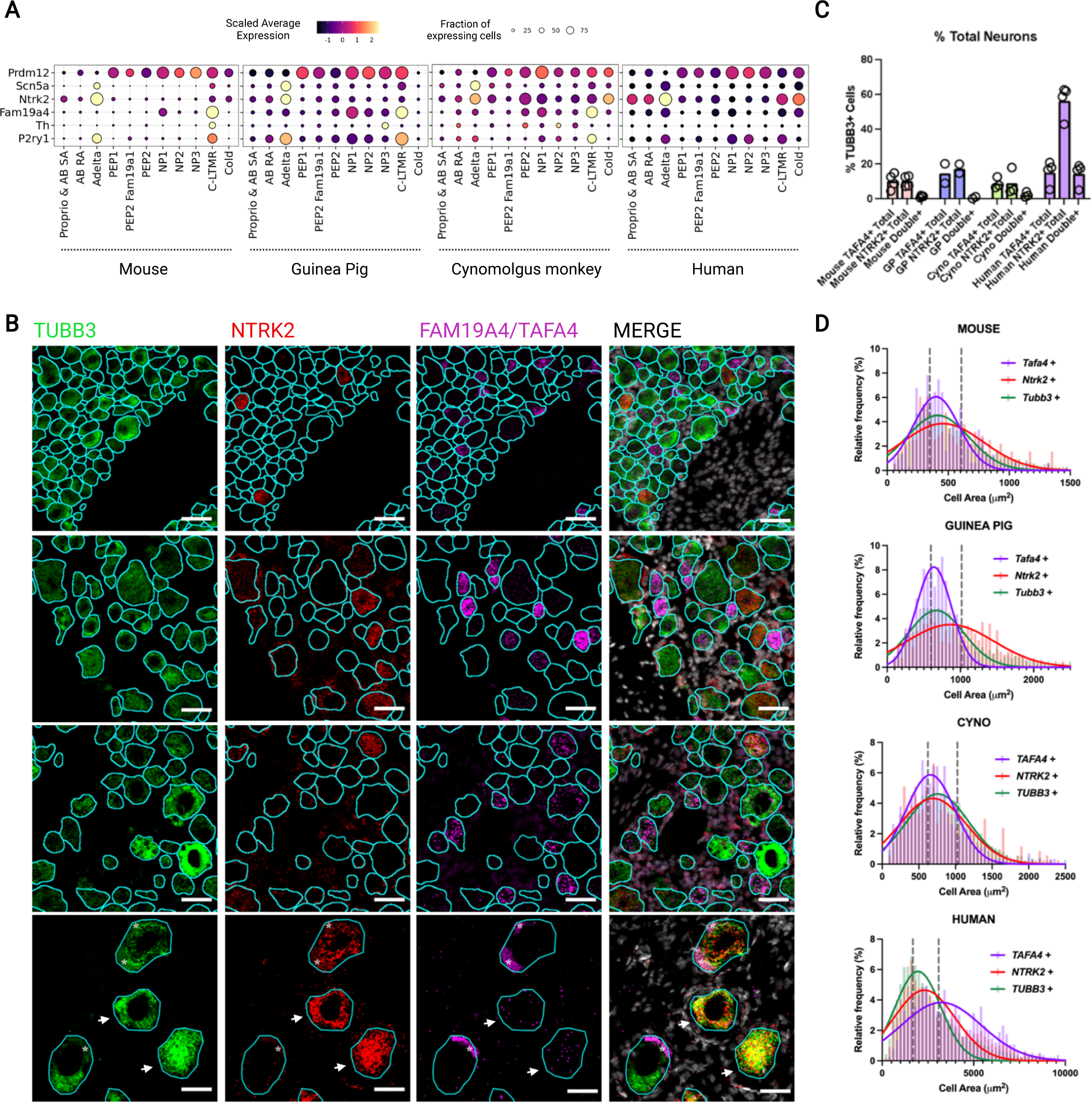
Divergent expression of TAFA4 across species. **a** Dot plots showing fraction of nuclei (size) and their scaled average expression level (color) in each neuron subtype (columns) across mouse, guinea pig, cynomolgus monkey, and human for genes (row names). **b** RNAScope validation of FAM19A4, NTRK2, and TUBB3 in DRGs. White arrows indicate neurons (TUBB3+) expressing both TAFA4 and NTRK2 in human. Asterisk in the human images represent lipofuscin aggregates. Scale bar represents 50µm. **c** Percentage of TAFA+, NTRK2+ and TAFA4+/NTRK2+ (‘double +’) in mouse, guinea pig, cynomolgus monkey and human. d Size distribution of TAFA4+, NTRK2+, TUBB3+ cells in each species. Grey dotted lines represent the 33% and 67% percentiles of the TUBB3 + distribution as a surrogate for small, medium, and large diameter neurons.

Taken together, this analysis highlights that while DRG subtypes are, in general, conserved from pre-clinical models to humans, significant differences exist between species.

Understanding the transcriptional and molecular differences across species will be critical to enabling a more complete understanding of pain and other pathophysiological processes, and are for the development of effective therapeutics. Importantly, these data amplify the need for accurate, detailed, and well-controlled cross-species atlases to more fully understand similarities and differences between preclinical and human sensory neuron function.

## DISCUSSION

Comparative cross-species analyses using atlases of single nuclei data is a powerful approach that can provide a high-level view of the cellular composition of tissues and a detailed map of the molecular landscape of these tissues. Such analyses can inform our understanding of biological mechanisms within tissues and can provide perspective on how these mechanisms might have evolved across species. Importantly, these data will play a central role in informing the translation of data from rodents to primate models and humans. Our integrative and comparative analysis of DRGs from mouse, guinea pig, cynomolgus monkey, and human identified diverse subtypes of DRG sensory neurons in each species, and showed that these subtypes are generally conserved. Our analysis highlighted conserved transcriptional features which may reflect core biological mechanisms that are relevant for subtype-specific sensory functions. Additionally, we identified divergent expression of molecular determinants involved in sensory function which highlight species-specific adaptations for pathophysiological processes like pain and tissue repair. Among those divergently expressed genes, we validated TAFA4, a known key modulator of the neuro-immune axis. TAFA4 is expressed in distinct populations of sensory neurons across mouse and human DRGs, which is relevant for therapeutic strategies targeting TAFA4. This paper provides a comprehensive, harmonized, and interpretable cross-species atlas and analysis of sensory neurons that will be crucial for better understanding of sensory neuron function and will enable the development of more effective pain and neuro-immune therapeutic targets.

Isolation of nuclei for the generation of single nuclei atlases can require extensive protocol development which is highly dependent on the type and state of the tissue sample ^32–34^. For our analysis of DRGs, we found that isolating nuclei from fresh DRG tissues using density-gradient centrifugation, as compared to FACS, enables improved capture of sensory neuron nuclei versus non-neuronal nuclei. We suspect that the reduced number of neuronal nuclei resulting from FACS could be due to the larger size of sensory neuron nuclei, making them more susceptible to the shear forces from the sorting process. Related, we note the possibility that some cell populations within the DRG may not be recovered due to the nature of either the cell isolation method or the droplet-based technology. For example, the microfluidic channel within a 10X Genomics Chromium chip is approximately 50-60 um in diameter which is not compatible with larger nuclei such as those from primate DRGs. Using our protocol, we also demonstrated the successful recovery of sensory neuron nuclei from frozen human tissues, which will expand future opportunities for utilizing archived clinical tissues to generate additional atlases of transcriptome data. Single nuclei preparation protocols will likely need to be optimized for different tissue types, sample conditions, or cell types (i.e., as non-neuronal cells present in the DRG). It is likely that our protocol will work well for generating atlases of other sensory neurons such as those from the nodose ganglia. Future analysis using high resolution spatial technologies such MERFISH^35^, or methods to characterize cell-type specific electrophysiological activity such as Patch-seq^36^, would provide a more complete picture of primate sensory DRG neurons.

To fully leverage these data, we explored different integrative computational approaches to align homologous sensory neuron subtypes across species and build a comprehensive cross-species atlas from preclinical models to humans. Some of these approaches included Seurat to aggregate datasets, and MetaNeighbor to identify cell types that are highly replicated among datasets. Each of these methods has distinct caveats. For example, the data aggregation algorithm implemented in Seurat can be susceptible to over-integration when only a subset of the cell populations is preserved among datasets. In addition, MetaNeighbor was developed to describe the extent of cell-type reproducibility across scRNA-seq data sets, rather than quantitatively classifying cell-types between query and reference data sets. However, in our analysis, the cell-type homology mapping results from Seurat and MetaNeighbor were well-aligned, suggesting that these results are robust and provide a computational framework for more systematically performing such cross-species comparison. Our computational work yielded a comprehensive cross-species atlas that described 11 transcriptionally distinct DRG sensory neuron subtypes across 4 species, using common subtype nomenclature. Continued work validating these computational findings through in-situ hybridization or other functional experiments will be essential to more precisely characterize these sensory neurons across species.

Although our comparative analysis using Seurat and MetaNeighbor revealed that DRG subtypes are, in general, well conserved across species, we performed additional analyses to assess transcriptional features reflecting core sensory neuron functions or species-specific adaptations. Interestingly, in the context of their gene expression profiles across species, we found that Proprioceptor & AB SA-LTMRs are the most conserved sensory neuron subtypes whereas NP1 and PEP2 Fam19a1 were the least conserved. This would be consistent with the important role that proprioceptors play in sensing space and limb positioning, or enabling fast muscle reflexes, all of which are crucial for vertebrate survival. Whereas sensations perceived by other sensory neuron subtypes are more tuned to an animal’s immediate environment and are manifest in gene expression variability within these subtypes. Differential expression analysis highlighted conserved gene expression programs that could reflect the distinct cell-type specific sensory function. In particular, the identification of conserved genes involved in pain signaling and acute inflammatory response among the peptidergic nociceptors present opportunistic therapeutic targets and inform our understanding of clinical programs targeting sensory neuron function. We also identified differences in the proportion of types of sensory neuron (i.e. C-LTMRs or NPs) across species. These differences in proportion could reflect differences in biology across the species or levels of DRGs. For example, differences in the C-LTMRs may reflect species-specific differences in the perception of specific cues, including those related to glabrous and hairy skin, or perception of itch. Alternatively, the differences in proportion may reflect that our single nuclei protocol did not capture these subtypes efficiently. It would be interesting to explore these differences in DRG subtype composition using subtype specific markers in tissue samples. Together, these data help identify different biological mechanisms that will ultimately inform therapeutic development targeting sensory neurons.

Although our data captured sensory neuron subtypes that are similarly described in recently published analysis of primate DRGs data^14–16^, we noted some significant differences in resolution of subtype classification and molecular marker expression which are relevant for interrogating sensory mechanisms from pre-clinical models to human. For example, the H10 cluster in *Ngyuen et al*. and pruritogen enriched receptor cluster in *Tavares-Ferreira et al.* map to both NP1 and NP2 in our data. Additionally, while *Tavares-Ferreira et al.* noted that they did not observe known C-LTMR-specific markers including TAFA4, P2RY1 and Zpf521/ZNF521 expression in their putative C-LTMR, we observed these markers’ expression in our human C-LTMR (Fig. 3d). These differences likely reflect a combination of factors including the number of cells sequenced, the resolution of the sequencing technology that was used, or the data integration framework used in the analysis, which can all have an impact on sensory neuron subtype annotation. It will be challenging to map therapeutically relevant markers from mouse to human subtypes with less resolved subtypes.

We used our atlas to investigate genes previously studied for their role in sensory function and pathophysiological processes to determine whether such genes might be functionally conserved across species. An intriguing example from this analysis was Tafa4/Fam19a4, a chemokine-like protein known to be secreted by IB4+ DRG neurons in rodents that has been studied for its role in chronic pain and tissue repair through the activation of the Lrp1 receptor on myeloid cells.

Within rodents we found that *Tafa4/Fam19a4* is predominantly expressed in C-fiber LTMRs and a subclass of NP nociceptors. In contrast, in humans *TAFA4/FAM19A4* was expressed in a broader set of sensory neurons including a subclass of A-fiber LTMRs and TRPM8+ Cold thermoreceptors, with limited expression detected in NP nociceptors and C-LTMRs. Consistent with our expression data, FISH analysis validated the differential expression of *TAFA4/FAM19A4* in human DRGs. It will be interesting to determine if the functional neuro-immune signaling axis mediated by TAFA4/FAM19A4-LRP1 is conserved in humans, even though Tafa4 may not be produced by the same subsets of sensory neurons. Additionally, our data also show that TRPM8+ sensory neurons also express TAFA4 in humans, while this is not observed in data from our rodent models. Interestingly, TRPM8 agonists have been used for pain inhibition in clinical settings ^37, 38^ and it could be that in humans TAFA4 and TRPM8 function together to mediate pain perception. Future work exploring the role of TAFA4/FAM19A4 in pain or tissue repair could provide insights that inform our efforts to translate therapeutic targets from animal models to the clinic.

The strength of the work presented here includes the development and use of a consistent laboratory protocol to avoid technical artifacts that can occur during single cell sequencing; generation of single nuclei data representing a broad sampling of sensory neurons across multiple species; application of multiple computational approaches to provide confident and detailed cell-type mappings across species; and harmonized nomenclature of sensory neuron subtypes from mouse, guinea pig, cynomolgus monkey, and human. Appropriate pairing of laboratory methodologies and computational approaches can have a significant impact on the development and ultimate downstream utility of cross-species atlases. Our ability to identify, characterize, and validate the differential expression of Tafa4 across species, highlights a significant example of the utility of our approach both in our ability to understand neuro-immune compartment and to develop clinically relevant therapeutics. Our analysis provides a more complete understanding of the general principles driving somatosensory mechanisms and informs our understanding of the role of sensory neurons in the maintenance of tissue homeostasis. Collectively, this work enables the development of more effective therapeutics targeting sensory neuron function.

## ACKNOWLEDGEMENTS

We thank Will Ewart, Orland Zuniga, Wyne Lee, and Aaron Fullerton for DRG tissues sourcing; C.K. Poon, Alice Tan, and Terence Ho for FACS support; Yuxin Liang, Qixin Bei, and Zora Modrusan for sequencing support; Bernard Chow, Aaron Lun, Altaf Kassam and Ramanandan Prabhakaran for reference genome support; Maggie Crow, Kevin Huang, Graham Heimberg, and Brad Friedman for helpful discussion on computational analysis; Jesse Hensen, Maggie Crow and Casper Hoogenraad for providing feedback on the manuscript. The graphical abstract was generated with BioRender. Funding was provided by Genentech.

## AUTHOR CONTRIBUTIONS

M.J., M.D., D.H.H., L.R-B., and J.S.K. designed the study with input from J.M. and A.J. M.D. performed protocol development and snRNA-seq data generation. M.J. and J.S.K. designed and performed computational analyses. J.M. and A.J. performed IHC validation and image analysis. M.D., M.B., and O.F. coordinated tissue sample procurement and processing. M.J., M.D., D.H.H., L.R-B., and J.S.K. wrote the manuscript with input from all authors.

## MATERIALS & CORRESPONDENCE

correspondence and requests for materials should be directed to Joshua S. Kaminker.

## METHODS

### EXPERIMENTAL MODEL AND SUBJECT DETAILS

#### Animals

Care and handling procedures of animals were reviewed and approved by the Genentech Institutional Animal Care and Use Committee (IACUC) and animal experiments were conducted in full compliance with IACUC policies and NIH guidelines.

Mice used in this study were C57BL/6J (The Jackson Laboratory, Stock No: 007914). Mice used in this study were 6-17 weeks old and both female and male mice were used. Hartley guinea pigs (Charles River Laboratory) used in this study were 5.5-6 months old and only female guinea pigs were used.

#### Tissues

Cynomolgus monkey DRGs from 3 animals were purchased from Covance in 2 shipments, Fresh (isolated, placed in Hibernate A solution (BrainBits, Catalog #HALF500), delivered at 4 degrees overnight) and Frozen (flash frozen at isolation, delivered on dry ice). The age of the animals varied from 8-9 years and only female monkeys were used.

Frozen Human DRGs were obtained from Anabios (6 donors) and Donor Network West (1 donor; 1 pair of lumbar 4 level). The DRG tissues from Anabios were stored in liquid nitrogen prior to shipment. All samples were assessed for tissue integrity prior to downstream applications.

Additional details on individual animals and samples are available in Supplementary Table 1.

### METHOD DETAILS

#### Tissue processing

Mice were euthanized by CO2 inhalation and decapitation. DRGs from lumbar 1-6 levels from both right and left sides were extracted, de-sheathed and placed in 1mL ice-cold lysis buffer (20 mM NaCl, 5 mM MgCL2, 0.1% TX-100, 10 mM Tris-HCl pH7.2) containing EDTA-Free protease inhibitor (Sigma, Catalog #4693124001), RNAse inhibitor 0.2 U/mL (Life Technologies, Catalog # N8080119), and RNase-free DNase (Promega Corporation, Catalog #M6101), in a 2 mL Dounce homogenizer tube (Kimble Chase, Catalog #885300-0002).

Guinea pigs were euthanized using protocols approved by the Genentech institutional Care and Use Committee. Briefly, the animals were placed in an isoflurane chamber for anesthesia. Once anesthetized, the animals were injected with a 1 mL solution of Euthasol and Sterile saline in a 1:1 ratio. Death was confirmed by loss of heartbeat and corneal opacity before decapitation and DRG dissection. DRGs from lumbar 3-5 levels from both right and left sides were collected and placed in the lysis buffer described above.

DRG samples from cynomolgus monkey and human were first cut into ∼1-2 mm pieces with a scalpel while placed on a dissection tray over dry ice and then placed in the lysis buffer described above.

#### Nuclei isolation from experimental animals

Dounce homogenization was used to dissociate all DRG tissues. Before douncing, 1 mL HBSS (ThermoFisher, Catalog #14025092) containing 3% BSA Fraction VI and RNAse inhibitor in nuclei suspension buffer (NSB) was added to the 2 mL Dounce Tissue Grinder (Kimble Chase, Catalog #885300-0002). The DRGs were homogenized with an A (“loose”) pestle using 5 to 10 strokes. The homogenate was then filtered through a 70-micron filter and spun down. The pellet was resuspended in the NSB. The frozen DRGs were not allowed to thaw before placing in the lysis buffer and douncing.

##### Density-gradient

The homogenate was layered over an Optiprep density (Sigma, Catalog #D-1556; 35%, 16%, 8% for rodents and 40%, 20% and 10% for primates) gradient and centrifuged at 2500 g for 20 min. Nuclei at the 16/35 interface were aspirated and mixed with an equal volume of NSB. Nuclei were re-pelleted, washed, and counted using a hemocytometer (gradient purified) prior to loading into a 10X Genomics Chromium controller.

##### FACS

Prior to sorting, nuclei from the density gradient were pelleted by centrifugation, labeled with propidium iodide (Life Technologies, Catalog #P1304MP) and DAPI (Life Technologies Catalog #62248) and sorted using a FACSAria Fusion Flow Cytometer (BD Biosciences)(Nuclei were selected based on double labeling with DAPI and PI and sorted into eppendorf tubes containing 0.5 mL nuclei suspension buffer (NSB, described above). Nuclei were then counted, pelleted and resuspended into an appropriate volume prior to loading into a 10X Genomics microfluidic chip for droplet generation and barcoding.

#### Preparation of single-nuclei RNA-sequencing libraries

Chromium Next GEM Single Cell 3ʹ GEM, Library & Gel Bead Kit v3.1 (10X Genomics PN-1000121) were used for library preparation according to the manufacturer’s user guides. The Cell-RT mix was prepared to aim for 10,000 nuclei per sample and applied to the Chromium^TM^ Controller for GEM generation and barcoding. Then samples were subjected to post GEM-RT cleanup, cDNA amplification (11 cycles with v3.1), and library construction according to the user manual. Sample index PCR was done with 12 cycles. Libraries were then quantified by Qubit dsDNA HS Assay Kit (Thermo Fisher Q33230) and profiled by Bioanalyzer High Sensitivity DNA kit (Agilent Technologies 5067-4626). Libraries were sequenced by HiSeq4000 (Illumina) following the 10X Genomics sequencing specification.

#### Mouse, guinea pig, cynomolgus monkey, and human reference genomes and transcriptomes

For genomic mapping, we augmented GRCm38, Cavpor3.0, macFas5, hg19 reference transcriptomes with introns to allow both pre-mRNAs and mature mRNAs to be mapped. GRCm38, Cavpor3.0, macFas5 and hg19 reference transcriptomes were modified according to the instructions provided by the 10X Genomics website (https://support.10xgenomics.com/single-cell-gene-expression/software/pipelines/latest/ advanced/references).

#### Single-nuclei data processing

##### Preprocessing and alignment

Single-cell RNA sequencing data were processed with a CellRanger analysis pipeline. Briefly, reads were demultiplexed based on perfect matches to expected cell barcodes. Transcript reads were aligned to the appropriate species genome using GSNAP (2013-10-10) (Wu and Nacu, 2010). Only uniquely mapping reads were considered for downstream analysis. Transcript counts for a given gene were based on the number of unique UMIs for reads (up to one mismatch). Both intronic and exonic reads were used to determine transcript count. Cell barcodes from empty droplets were filtered by requiring a minimum number of detected transcripts. Data quality for individual libraries was assessed based on total read depth, percentage of reads with valid barcodes, percentage of demultiplexed reads in detected cells, number of detected cells, and number of analyzed cells. Sample quality was further assessed based on the distribution of per-cell statistics, such as total number of reads, percentage of reads mapping uniquely to the reference genome, percentage of mapped reads overlapping exons, number of detected transcripts (UMIs), number of detected genes, and percentage of mitochondrial transcripts.

##### Removal of background noise in gene expression matrices

We used the ‘remove-background’ function of CellBender (v.0.2.0) to remove technical ambient RNA counts and empty droplets from the gene expression matrices^39^. Cell Ranger-generated ‘raw_feature_ bc_matrix.h5’ files were used as input for CellBender. The parameter ‘expected-cells’ was obtained from the Cell Ranger metric ‘Estimated Number of Cells’, while the parameter ‘total-droplets-included’ was set to a value between 8,000 and 12,000 to represent a point within the plateau of the barcode rank plot in all samples.

##### Quality control and clustering

After this initial quality control, nuclei with less than 1,000 total UMIs, 500 unique detected genes and greater than 25% mitochondrial UMIs were discarded. After the filtering step, the gene x cell matrix of raw UMI counts was log-normalized using ‘NormalizeData()’ in SeuratV3 (Stuart et al., 2019). All libraries within each species were integrated using ‘FindIntegrationAnchors()’ and ‘IntegrateData()’ functions in SeuratV3. Then, we scaled the species-specific integrated data, performed dimensionality reduction by PCA, calculated UMAP coordinates and Louvain clustering for all nuclei using SeuratV3 (Fig. 2a-b, 3a-c, Supplementary Fig. 3a-c).

| Dataset | Cell type | Used PCs |
| --- | --- | --- |
| Mouse DRG | All cell types | 30 |
| Mouse DRG | Neurons | 30 |
| Guinea Pig DRG | All cell types | 30 |
| Guinea Pig DRG | Neurons | 30 |
| C. Monkey DRG | All cell types | 30 |
| C. Monkey DRG | Neurons | 30 |
| Human DRG | All cell types | 30 |
| Human DRG | Neurons | 30 |

Major cell type clusters were identified based on top differentially expressed genes from each cluster (Fig. 2a-b,3a-c, Supplementary Fig. 3a-c). In all species-specific data, clustering was performed twice: first, to separate neurons and glia from other cells, and then to sub-cluster the DRG neurons to obtain high-resolution clusters within the DRG neuron group.

#### DRG neuron signature score

The DRG neuron signature score reflects the mean expression levels of a set of marker genes for neurons. We compiled the following lists of neuronal marker genes for DRG neurons from the literature^7, 14, 16^: Ret, Ldhd, Nefh, Cntnap2, Scn9a, Scn8a, Scn10a, Scn11a, Tac1, Plxnc1, Gfra2, Calca, Slc17a7, S100b, Piezo2, Uchl1, Rbfox3, Snap25. We calculated the average expression of these genes to construct the “DRG neuron” signature score (Fig. 1f), and used this score to assess neuron nucleus quality across nucleus isolation methods and tissue types.

#### Differential expression analysis between isolation methods and sample types

To compare the quality of transcriptomic profiles generated by different isolation methods and sample types, we performed differential expression (DE) analysis. Pseudo-bulk expression profiles were derived from single-cell datasets by calculating the average of the total number of UMIs for each gene across all nuclei of each sample. This gave a gene-by-pseudobulk count matrix which was then normalized to a normalized count statistic using the ‘calcNormFactors()’ function from edgeR. DE analysis was performed by calling ‘glmLRT()’ and then using ‘topTags()’ to extract the final differential expression statistics.

#### DRG neuron subset specific analysis

To obtain high-resolution clusters within the DRG neuron subset in all species-specific data, we first removed all non-neuronal nuclei barcodes and then nuclei that express any satellite glial specific transcripts (PLP1 < 1 & MPZ < 1 & SPARC < 1) were removed. The resulting digital gene-expression matrix (DGE) was carried forward for clustering.

We annotated different subsets of large diameter myelinated A-LTMRs using Nefh, Pvalb, Spp1, Calb1, Ntrk3, Scn5A, Ntrk2, Necab2, Cntnap2, and Fam19a1. Non-peptidergic C-fiber nociceptors(NPs) subsets were annotated using Gfra1, Gfra2, Trpc3, Lpar3, Mrgpra3, Mrgprd, Sst, Il31ra, Nppb, Trpv1, Trpa1, Ret, Scn10a, Scn11a, P2rx3, and Plxnc1. C-fiber peptidergic nociceptors (PEPs) subsets were annotated using Tac1, Adcyap1, Gal, Kit, Calca, Ntrk1, Trpa1, Scn10a and Scn11a. Cold thermoreceptors subsets were annotated using Trpm8, Tac1, Foxp2, Cdh8, Penk1, and Piezo2. Finally, C-LTMRs were annotated using Th, Slc17a8, Fam19a4, Gfra2, Piezo2 and Zfp521/ZNF521.

To avoid having one species dominate the downstream analyses including integration and to account for potential differences in each species’ clustering resolution, we downsampled the number of nuclei to have similar numbers across species at each DRG subtype cluster (e.g., A-LTMRs, PEPs, NPs,C-LTMRs) using the ‘downsample’ argument in the ‘subset()’ function of SeuratV3. These downsampled DGEs were used for cross-species cell-type mapping analyses including MetaNeighbor and Integration.

#### MetaNeighbor analysis

MetaNeighbour v1.9.1 (RRID SCR_016727) was used to provide a measure of neuronal subclass and cluster replicability within and across species (Scripts and tutorials are available on GitHub (http://github.com/gillislab/MetaNeighbor)). The mean area under the receiver operator characteristic curve (AUROC) scores from MetaNeighbor were used as a proxy for the quantitative similarity between cell-type pairs. We performed two rounds of MetaNeighbor analysis, first on the combined all species (mouse, guinea pig, cynomolgus monkey and human)-all cell types dataset and all species-DRG neuron-specific subsets(downsampled).

Highly variable genes were identified using the ‘get_variable_genes()’ function, yielding 493 genes for all cell types dataset and 390 genes for the DRG neuron subset. These were used as input for the ‘MetaNeighbourUS()’ function, which was run using the fast_version. AUROCs are plotted in heat maps in Fig. 3c and Supplementary Fig. 3c. In all species-all cell types MetaNeighbor analysis, the dendrogram of AUROC scores were organized according to major cell-types rather than species, suggesting that cell type similarity transcends mammalian species differences (Supplementary Fig. 3c).

#### Cross-species dataset integration analysis

To identify homologous cell types across species, we used Seurat’s CCA workflow to perform a separate supervised integration of DRG neurons across species. Downsampled raw expression matrices were reduced to include only those genes with one-to-one orthologues defined in the three species (downloaded from http://www.ensembl.org/biomart/martview) and placed into Seurat objects with accompanying metadata. To integrate across species, all Seurat objects were merged and normalized using SeuratV3. We explored both rPCA and CCA for integration and these methods produced similar results (data not shown).

#### Identification of core and species-specific transcription profiles for major DRG neuron classes

To identify conserved and species-specific transcriptional signatures for each neuron class (i.e., A-LTMRs, PEPs, NPs,C-LTMRs, Cold thermoreceptors), we used expression matrices that were reduced to include only those genes with one-to-one orthologues. Within each species, we performed differential expression(DE) analysis using ‘FindAllMarkers()’ function in SeuratV3 to identify top markers in each neuron subtype/class. While using ‘FindAllMarkers()’ function, we set ‘logfc.threshold’ = 0.5 and ‘min.pct’ = 0.3 for requiring top marker genes to be present in > 30% of nuclei and on average, have a log2 fold-difference > 0.5 between two testing groups.

We then intersected these DE lists to identify DE orthologs that were either shared by all species (e.g., conserved A-LTMRs, conserved NPs, conserved PEPs, conserved C-LTMRs) or unique to each species (e.g., mouse A-LTMRs, guinea pig A-LTMRs, cyno. monkey A-LTMRs, human A-LTMRs), within each neuron class (Fig. 4c).

#### Transcriptome correlation analysis

To investigate transcriptome similarity from mouse to human, we performed correlation analysis on gene expression data. Within each species, we first summed all transcript counts for each DRG neuron subtype and log-transformed the data with the ‘log2()’ function in R. Then we calculated spearman correlation using the ‘cor()’ function with the ‘method’ argument set to ‘spearman. The correlation coefficients were plotted for different human-other species pairs (Fig 4e).

#### Gene Ontology analysis

Gene ontology enrichment analysis was performed using the ‘enrichGO()’ function from the clusterProfiler R package in which p-values are calculated based on the hypergeometric distribution and corrected for testing of multiple biological process GO terms using the Benjamini-Hochberg procedure ^40^. GO terms were accessed using the AnnotationHub R package.

#### Fluorescent *in situ* hybridization (RNAScope)

DRGs from mouse and guinea pig were obtained as described above. The nerves and connective tissues were trimmed, and DRGs were placed in OCT molds and frozen rapidly in a mixture of dry ice and Ethanol. 5-10 micron sections were cut using a cryostat and placed on slides for downstream RNA labeling studies.

RNAScope Multiplex Fluorescent Kit (Advanced Cell Diagnostics) was used per manufacturer’s recommendations for fresh-frozen samples with the following alterations. During pretreatment, sections were treated with hydrogen peroxide for 10min. And Protease IV for 20min. prior to the addition of relevant probes. Opal dyes 690, 570, and 520 (Akoya Biosciences) were used for fluorescence and after the final HRP block step, samples were stained with DAPI solution for 1min. followed by mounting with ProLong Gold Antifade Mounting Solution (Thermo Fisher Scientific, Cat#P36961). Probes used for RNAscope (Advanced Cell Diagnostics) include: For mouse: Rgs5 (Cat #430181), Th (Cat #317621), Slc17a8 (Cat #431261), Tafa4 (Cat #813621), Ntrk2 (Cat #423611), Tubb3 (Cat #423391). For guinea pig: Tafa4 (custom, Cat #1128881), Ntrk2 (custom, Cat #1128891), mouse Tubb3 used for pan-neuronal marker. For cynomolgus monkey: Tafa4 (custom, Cat#1128871), Ntrk2 (Cat#424151), used human Snap25 probe for pan-neuronal marker. For human: Tafa4 (custom, Cat#1037841), Ntrk2 (Cat#402621), Snap25 (Cat#518851).

#### Fluorescence microscopy and image analysis

Epifluorescence images were taken using a Zeiss Axio Imager.M2 upright microscope equipped with an Apotome.2 structured illumination module. Images were acquired using a Zeiss Colibri 7 LED, DAPI/AF488/AF555/Cy5 filter sets, a Plan-Apochromat 20X/0.8 objective lens, and a Hamamatsu ORCA-Flash 4.0 Digital CMOS camera. 5 z-stack images were collected at 2um intervals and maximum intensity projections were created with Zeiss Zen software. For comparison of mouse C-LTMR1 and C-LTMR2 populations, all images were acquired using the same LED intensity and exposure time settings. For cross-species NTRK2/FAM19A4/TUBB3 imaging, the image acquisition settings were the same for the set of images within a given species.

All image analysis was performed in ImageJ/FIJI using custom macros. For comparison of mouse C-LTMR1 and C-LTMR2 subtypes, manual regions-of-interest (ROIs) were drawn using the *Slc17a8* signal to mark the entire population of putative C-LTMRs in each section. The optical density of the *Slc17a8*, *Rgs5*, and *Th* channels were measured within each ROI as well as the average optical density of putative single transcripts, identified as distinct, round puncta with clearly decaying intensity on all sides, for each channel per section. Following background correction, the number of puncta per ROI was calculated by dividing the optical density of a given ROI by the average optical density of a single puncta. All measurements were normalized to the cross-sectional area of a given ROI and are reported as the number of puncta per um^2. We manually determined a threshold for *Th* high and low cells (dotted line in Fig. 2f) and separated the cells into putative C-LTMR1 and C-LTMR2 populations, respectively, based on the snRNAseq analysis. 3 DRG sections per mouse were imaged per vertebral level (cervical, thoracic, lumbar) and the values were averaged across technical replicates; data in Fig. 2f represents each independent biological replicate. DRGs from 2 male and 2 female mice were analyzed and we did not detect any difference between sexes therefore the data was combined. Data in Fig. 2g is combined for all *Slc17a8*-defined ROIs across all replicates for each vertebral level.

For the cross-species comparison of *NTRK2*/*FAM19A4*/*TUBB3* expression, neuronal ROIs were segmented automatically with the Cellpose algorithm ^41^ for mouse, guinea pig, and cynomolgus macaque using the *TUBB3* images. For human, neuronal ROIs were drawn manually using the *TUBB3* and DAPI signals due to interference of pervasive autofluorescence/lipofuscin that appeared as diffuse signal across the AF488/AF55/Cy5 channels, even in the negative control probe images. For mouse, guinea pig, and cynomolgus macaque, a similar analysis pipeline as described above was applied where the optical density for each channel within the neuronal ROIs was measured along with the average optical density of putative single transcripts for each probe. The number of puncta was normalized to the cross-sectional area per neuronal ROI and a threshold for *NTRK2*-positive and *FAM19A4*-positive cells was manually determined for each species. For human images, thresholds for *NTRK2*-positive and *FAM19A4*-positive cells were manually defined based on exceeding any autofluorescence and background signal observed in negative control images. Each individual neuronal ROI was manually defined as *NTRK2*-positive, *FAM19A4*-positive, or double-positive. Data in Fig. 5c was collected from lumbar DRGs and averaged across sections per biological sample (i.e. animal): 3 sections per mouse from 2 male and 2 female mice, 3 sections per guinea pig from 2 female guinea pigs, 2 sections per cynomolgus macaque from 3 female macaques, and 2-3 sections per human from 2 male and 2 female humans. Histograms in Fig. 5d were generated using cross-sectional area data from all *NTRK2*-positive, *TAFA4*-positive, and *TUBB3*-positive ROIs for each respective species. Relative frequency histograms were generated and Gaussian curves were fit using GraphPad Prism v9. Gray dotted lines in each histogram represent the 33rd and 67th percentiles of the entire TUBB3 distribution for each species as an approximation of small, medium, and large neurons in each species.

All representative images are pseudo-colored with the brightness and contrast adjusted to improve visualization. C-LTMR or neuronal ROIs are overlayed in cyan. For human DRG images in Fig. 5b, autofluorescence/lipofuscin signals are denoted by white asterisks in the 3 individual channels and the merged image.

#### Data availability

Datasets have been deposited in the Gene Expression Omnibus (GEO) repository, accession number GSE201654. Datasets will be released to public when the manuscript is published on a peer-reviewed journal.

## REFERENCES

1. Abraira, V. E. & Ginty, D. D. The Sensory *Neuron*s of Touch. Neuron 79, 618–639 (2013).

2. Lumpkin, E. A. & Caterina, M. J. Mechanisms of sensory transduction in the skin. Nature 445, 858–865 (2007).

3. Lewin, G. R. & Moshourab, R. Mechanosensation and pain. *J Neurobiol* **61**, 30–44 (2004).

4. Moraes, E. R. de, Kushmerick, C. & Naves, L. A. Morphological and functional diversity of first-order somatosensory neurons. Biophysical Rev 9, 847–856 (2017).

5. Emery, E. C. & Ernfors, P. The Oxford Handbook of the Neurobiology of Pain. 127–155 (2018) doi:10.1093/oxfordhb/9780190860509.013.4.

6. Li, C.-L. et al. Somatosensory neuron types identified by high-coverage single-cell RNA-sequencing and functional heterogeneity. Cell Res 26, 83–102 (2016).

7. Renthal, W. et al. Transcriptional Reprogramming of Distinct Peripheral Sensory *Neuron* Subtypes after Axonal Injury. Neuron 108, 128–144.e9 (2020).

8. Usoskin, D. et al. Unbiased classification of sensory neuron types by large-scale single-cell RNA sequencing. Nat Neurosci 18, 145–153 (2015).

9. Rostock, C., Schrenk-Siemens, K., Pohle, J. & Siemens, J. Human vs. Mouse Nociceptors – Similarities and Differences. Neuroscience 387, 13–27 (2017).

10. Li, Y. et al. DRG Voltage-Gated Sodium Channel 1.7 Is Upregulated in Paclitaxel-Induced Neuropathy in Rats and in Humans with Neuropathic *Pain*. J Neurosci 38, 1124–1136 (2018).

11. Coward, K., et al. Immunolocalization of SNS&sol;PN3 and NaN&sol;SNS2 sodium channels in human pain states. Pain 85, 41–50 (2000).

12. Coward, K. et al. Plasticity of TTX-sensitive sodium channels PN1 and Brain III in injured human nerves. Neuroreport (2001) doi:10.1097/00001756-200103050-00014.

13. Fukuoka, T. et al. Comparative study of the distribution of the α-subunits of voltage-gated sodium channels in normal and axotomized rat dorsal root ganglion neurons. J Comp Neurol 510, 188–206 (2008).

14. Nguyen, M. Q., Buchholtz, L. J. von, Reker, A. N., Ryba, N. J. & Davidson, S. Single-nucleus transcriptomic analysis of human dorsal root ganglion neurons. Elife 10, e71752 (2021).

15. Tavares-Ferreira, D. et al. Spatial transcriptomics of dorsal root ganglia identifies molecular signatures of human nociceptors. Sci Transl Med 14, eabj8186 (2022).

16. Kupari, J. et al. Single cell transcriptomics of primate sensory neurons identifies cell types associated with chronic pain. Nat Commun 12, 1510 (2021).

17. Stuart, T. et al. Comprehensive Integration of Single-Cell Data. Cell 177, 1888–1902.e21 (2019).

18. Crow, M., Paul, A., Ballouz, S., Huang, Z. J. & Gillis, J. Characterizing the replicability of cell types defined by single cell RNA-sequencing data using MetaNeighbor. Nat Commun 9, 884 (2018).

19. Hodge, R. D. et al. Conserved cell types with divergent features in human versus mouse cortex. Nature 573, 61–68 (2019).

20. Bakken, T. E. et al. Comparative cellular analysis of motor cortex in human, marmoset and mouse. Nature 598, 111–119 (2021).

21. Wang, J. et al. Tracing cell-type evolution by cross-species comparison of cell atlases. Cell Reports 34, 108803 (2021).

22. Chiu, I. M. et al. Bacteria activate sensory neurons that modulate pain and inflammation. Nature 501, 52–57 (2013).

23. Oetjen, L. K. et al. Sensory Neurons Co-opt Classical Immune Signaling Pathways to Mediate Chronic Itch. Cell 171, 217–228.e13 (2017).

24. Cohen, J. A. et al. Cutaneous TRPV1+ Neurons Trigger Protective Innate Type 17 Anticipatory Immunity. Cell 178, 919–932.e14 (2019).

25. Riol-Blanco, L. et al. Nociceptive Sensory Neurons Drive Interleukin-23 Mediated Psoriasiform Skin Inflammation. Nature 510, 157–161 (2014).

26. Russa, F. L. et al. Disruption of the Sensory System Affects Sterile Cutaneous Inflammation In Vivo. J Invest Dermatol 139, 1936–1945.e3 (2019).

27. Hoeffel, G. et al. Sensory neuron-derived TAFA4 promotes macrophage tissue repair functions. Nature 594, 94–99 (2021).

28. Yoo, S. et al. TAFA4 relieves injury-induced mechanical hypersensitivity through LDL receptors and modulation of spinal A-type K+ current. Cell Reports 37, 109884 (2021).

29. Yu, X. et al. Dorsal root ganglion macrophages contribute to both the initiation and persistence of neuropathic pain. Nat Commun 11, 264 (2020).

30. Kambrun, C. et al. TAFA4 Reverses Mechanical Allodynia through Activation of GABAergic Transmission and Microglial Process Retraction. Cell Reports 22, 2886–2897 (2018).

31. Li, L. et al. The Functional Organization of Cutaneous Low-Threshold Mechanosensory Neurons. Cell 147, 1615–1627 (2011).

32. Drokhlyansky, E. et al. The Human and Mouse Enteric Nervous System at Single-Cell Resolution. Cell (2020) doi:10.1016/j.cell.2020.08.003.

33. Slyper, M. et al. A single-cell and single-nucleus RNA-Seq toolbox for fresh and frozen human tumors. Nat Med 26, 792–802 (2020).

34. Habib, N. et al. Massively parallel single-nucleus RNA-seq with DroNc-seq. Nat Methods 14, 955–958 (2017).

35. Xia, C., Fan, J., Emanuel, G., Hao, J. & Zhuang, X. Spatial transcriptome profiling by MERFISH reveals subcellular RNA compartmentalization and cell cycle-dependent gene expression. P Natl Acad Sci Usa 116, 19490–19499 (2019).

36. Cadwell, C. R. et al. Electrophysiological, transcriptomic and morphologic profiling of single neurons using Patch-seq. Nat Biotechnol 34, 199–203 (2016).

37. Weyer, A. D. & Lehto, S. G. Development of TRPM8 Antagonists to Treat Chronic Pain and Migraine. Pharm 10, 37 (2017).

38. Fernández-Peña, C. & Viana, F. Targeting TRPM8 for Pain Relief. Open Pain J 6, 154–164 (2013).

39. Fleming, S. J., Marioni, J. C. & Babadi, M. CellBender remove-background: a deep generative model for unsupervised removal of background noise from scRNA-seq datasets. Biorxiv 791699 (2019) doi:10.1101/791699.

40. Yu, G., Wang, L.-G., Han, Y. & He, Q.-Y. clusterProfiler: an R Package for Comparing Biological Themes Among Gene Clusters. Omics J Integr Biology 16, 284–287 (2012).

41. Stringer, C., Wang, T., Michaelos, M. & Pachitariu, M. Cellpose: a generalist algorithm for cellular segmentation. Nat Methods 18, 100–106 (2021).

